# Monozygotic twin pairs discordant for amyotrophic lateral sclerosis carry both common and unique epigenetic differences relevant to disease

**DOI:** 10.1101/090977

**Authors:** Paul E Young, Stephen Kum Jew, Michael E Buckland, Roger Pamphlett, Catherine M Suter

## Abstract

Amyotrophic lateral sclerosis (ALS) is a devastating late-onset neurodegenerative disorder in which only a small proportion of patients carry an identifiable causative genetic lesion. Despite high heritability estimates, a genetic etiology for most sporadic ALS remains elusive. Here we report the epigenetic profiling of five monozygotic twin pairs discordant for ALS in whom previous genome sequencing excluded a genetic basis for their disease discordance. By studying cytosine methylation patterns in peripheral blood DNA we identified thousands of large between-twin differences at individual CpGs. While the specific sites of difference were largely idiosyncratic to a twin pair, a proportion (involving GABA signalling) were common to all affected individuals. In both instances the differences occurred within genes and pathways related to neurobiological function and dysfunction. Our findings reveal widespread changes in epigenetic marks in ALS patients, consistent with an epigenetic contribution to disease. These findings may be exploited to develop blood-based biomarkers of ALS and develop further insight into disease pathogenesis. We expect that our findings will provide a useful point of reference for further large scale studies of sporadic ALS.

## Non-Technical Summary

Amyotrophic lateral sclerosis (ALS) is a late-onset and fatal disease characterised by progressive loss of motor neurons and consequent loss of motor function. While about 10% of ALS cases are due to an inherited mutation in certain genes, about 90% are sporadic and most of these have no identifiable genetic cause. Here we looked for potential epigenetic changes associated with sporadic ALS by studying five sets of identical twins where only one twin was affected by ALS. By comparing DNA methylation patterns between affected and genetically-identical unaffected co-twins we identified thousands of epigenetic differences associated with ALS. Many of these changes occurred at genes with known neurological functions, implying that an epigenetic signature of ALS can be identified in peripheral blood. The epigenetic changes we have identified may prove to be useful biomarkers of disease and provide further insight into the underlying cause of sporadic ALS.

## Introduction

Amyotrophic lateral sclerosis (ALS), also known as motor neuron disease, is a lethal adult-onset disease that causes progressive muscle weakness, with death usually 2 to 5 years after initial diagnosis [1]. About 10% of ALS is familial and attributable to germline mutation of specific genes, but in the majority of cases (~90%) no other family member is affected, and the cause of most of this so-called sporadic form of ALS (SALS) remains unknown. Genetic, epigenetic and environmental factors have all been suggested to play a role in SALS, with combinations of these factors proposed to contribute to a multi-staged etiology [2].

Although rare single or multiple genetic variants may underlie some cases of SALS [3, 4], much of the heritability of the disease remains to be found [5]. Attention has turned to the possibility that epigenetic factors could contribute to ALS and its associated condition, frontotemporal dementia [6]. The fact that epigenetic changes may be therapeutically modified has driven research in this area [7]. A limited number of unvalidated epigenetic studies of SALS have been undertaken, involving single genes such as *SOD1* and *VEGF*[8], small groups of genes such as those in the metallothionein family (involved in detoxifying heavy metals) [9], and genome-wide methylation analysis using microarray [10]. However, the role of epigenetic variants in SALS remains unclear and largely unexplored [11].

Assessing the epigenetic basis of any disease in outbred populations such as humans is difficult since benign genetic variation is a major confounder [12]. Furthermore, there is the additional challenge of distinguishing germline epigenetic abnormalities from somatic changes secondary to either pre- or post-natal environmental influences [13]. This is particularly relevant to standard case-control studies because a vast number of environmental influences come into play within a normal human lifetime. One way of addressing this variability between subjects is to study disease-discordant monozygotic twins, who share at least the same genome, are exposed to a parallel intrauterine environment, and often have similar lifestyles. This is an appealing approach for ALS since twin registry studies show ALS is discordant in over 90% of monozygotic twins [14–16], which implies a major epigenetic or environmental component in disease susceptibility. Epigenetic differences certainly exist between monozygotic twins [17], and attempts have been made to link such co-twin differences to disorders as diverse as psoriasis [18], neurofibromatosis [19], and frontometaphyseal dysplasia [20].

In this study we explored the nature and extent of epigenetic changes in peripheral blood DNA from five sets of well-characterised ALS-discordant monozygotic twins. We compared genomic DNA methylation patterns between these twins in both case-control and co-twin analyses. We found a large number of differentially methylated sites between twins, most of which occurred at isolated CpGs, which cluster in common genes and pathways relating to neurobiological functions.

## Results

### Monozygotic twins discordant for ALS show no evidence of germline epimutation at known ALS genes

Ten individuals were included in this study: five individuals with a diagnosis of sporadic ALS, and their respective unaffected monozygotic twin siblings (**Table 1**). The average difference in time between ALS onset in the affected twin and the current age of the unaffected twin was 8.4 years (range 7–10 years), implying a non-genetic etiology of ALS in the affected twin. Consistent with this, none of the twins harboured an expanded repeat at the *C9orf72* locus [21]. Furthermore, previous whole genome sequencing failed to detect any other significant genetic variation between these co-twins; no pathogenic point mutation, insertion/deletion, or structural alteration was identified in the affected twins when compared with their unaffected co-twin [22]. We therefore considered the possibility that the underlying predisposing defect in the affected twins may be epigenetic in nature: epigenetic differences are not uncommon between monozygotic twins, and available evidence suggests that many such differences may be present from birth [17]. We obtained representative cytosine methylation profiles on peripheral blood DNA for each individual using Illumina 450K Infinium methylation arrays [23] and reduced representation bisulfite sequencing (RRBS) [24]. The 450K array assesses methylation at a predetermined set of ~450,000 single CpG sites concentrated around gene promoters and gene 100 bodies. RRBS assesses ~1% of the genome and ~1 million or more CpGs; it is complementary to 101 the 450K array since it also captures many CpGs outside of CpG islands and allows allelic 102 resolution of methylation patterns. In our RRBS libraries we obtained >10x coverage on an 103 average of 2.2 million CpGs and ≥20x coverage on an average of 1.4 million CpGs for each sample. Statistics on each of the twin RRBS libraries can be found in **Table S1**.

**Table 1.** Characteristics of ALS 106 and NonALS Monozygotic Twins

| Twins<br>(ALS/nonALS) | Gender | Age | Diagnosis<br>of affected<br>twin | ALS<br>discordance<br>(years) | ALS twin | NonALS twin |
| --- | --- | --- | --- | --- | --- | --- |
| Pair 1<br>(A/B) | Male | 52 | PMA<sup>§</sup> | 10 | Non-smoker | Ex-smoker |
| Pair 2<br>(C/D) | Male | 60 | ALS | 9 | Non-smoker,<br>depression | Non-smoker, boat-<br>building chemicals |
| Pair 3<br>(E/F) | Female | 53 | ALS | 8 | Ex-smoker, asthma | Ex-smoker, GORD,<br>statin |
| Pair 4<br>(G/H) | Female | 62 | ALS<sup>§</sup> | 8 | Ex-smoker, textile<br>chemicals | Smoker, nephritis,<br>COPD, breast<br>cancer, NHL |
| Pair 5<br>(I/J)* | Female | 69 | ALS | 7 | Non-smoker,<br>stroke, hepatitis A,<br>alcohol | Non-smoker, breast<br>cancer |

We first sought evidence for aberrant methylation in the affected twins at promoters of the ALS-associated genes *ALS2*, *C9orf72*, *FUS*, *OPTN*, *PFN1*, *SETX*, *SOD1*, *SPG11*, *TARDBP*, *VAPB*, *VCP* and *UBQLN2* [25]. We found no evidence of hypermethylation at probes within any known ALS gene promoter in any twin in the 450K array data. Further, at 10x coverage our RRBS libraries captured allelic information on the promoters of the same genes in all twin pairs, but none of the affected twins exhibited aberrant methylation at any of these loci (**Fig 1A**). Patterns of methylation at each known ALS disease locus were almost identical among all individuals, with all autosomal promoters showing little to no methylation, as shown for example in *C9orf72* (**Fig 1B)**. Thus, the discordance for ALS in these monozygotic twin pairs is not due to a germline genetic or epigenetic defect in any of the genes commonly associated with familial ALS.

**Fig 1.**
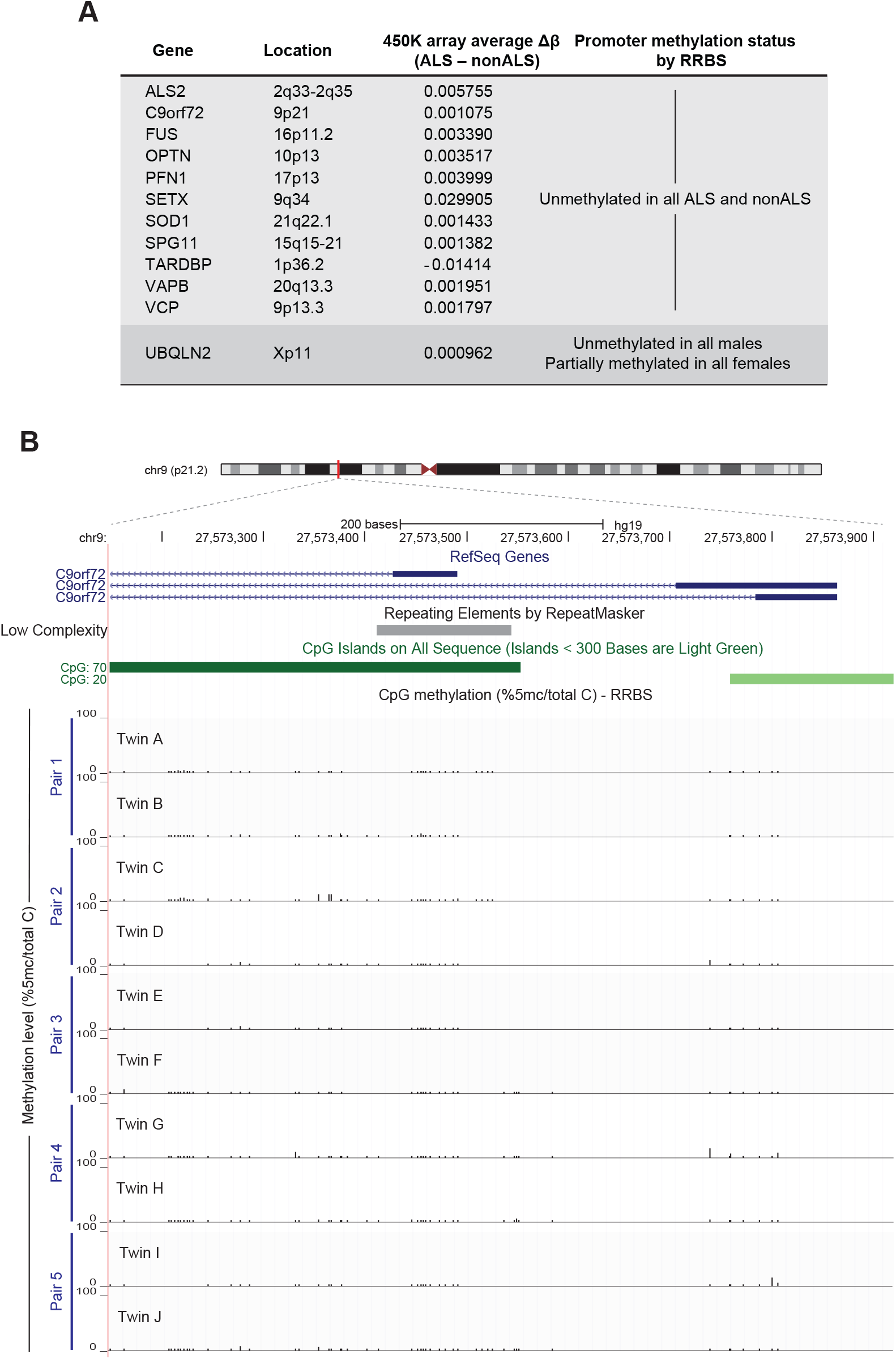
**DNA methylation patterns at known ALS gene promoters do not differ between monozygotic twins discordant for ALS (A)** Known ALS disease genes captured by both Illumina Infinium 450K array and RRBS at 10x coverage; Δβ represents the average difference in methylation levels between affected and unaffected twins. **(B)** Genome browser snapshot showing one representative example (the CpG island of *C9orf72*) of methylation patterns obtained by RRBS. The region harbouring the hexanucleotide repeat is shown by the grey bar under the RepeatMasker track. None of the twins harbour an expanded repeat, nor do they harbour significant methylation at any CpG across the *C9orf72* promoter.

### Case-control analysis of methylation implicates GABA receptor signalling as a commonly perturbed epigenetic network in ALS

We next took an unbiased approach to determining whether epigenetic differences may underlie the twin discordance for ALS. Unsupervised hierarchical clustering of RRBS data at 10x did not separate cases and controls, but instead identified five distinct clusters representing the five twin pairs (**Fig 2A**). This is not surprising given the known influence of genotype on inherited methylation patterns [26]. We then used the statistical package methylKit [27] to ask whether there were any differentially methylated CpG sites (DMCs) in common between all ALS cases versus all unaffected controls. At a significance threshold of *q*<0.01, this identified 135 CpG sites with ≥20% average difference in methylation between the two groups (**Fig 2B; Table S2**). About one half of these DMCs were in unannotated, intergenic regions of the genome, with the remainder predominantly within intronic regions (**Fig 2C**). Unsupervised clustering of the 450K data led to a similar clustering by twin pair, not disease status (**Fig 2D**). Analysis of the array data using minfi[28] failed to identify any significant common DMCs; CpGs with nominal significance, or approaching significance after correction for multiple testing, exhibited only tiny differences in methylation between cases and controls (**Fig 2E**).

**Fig 2.**
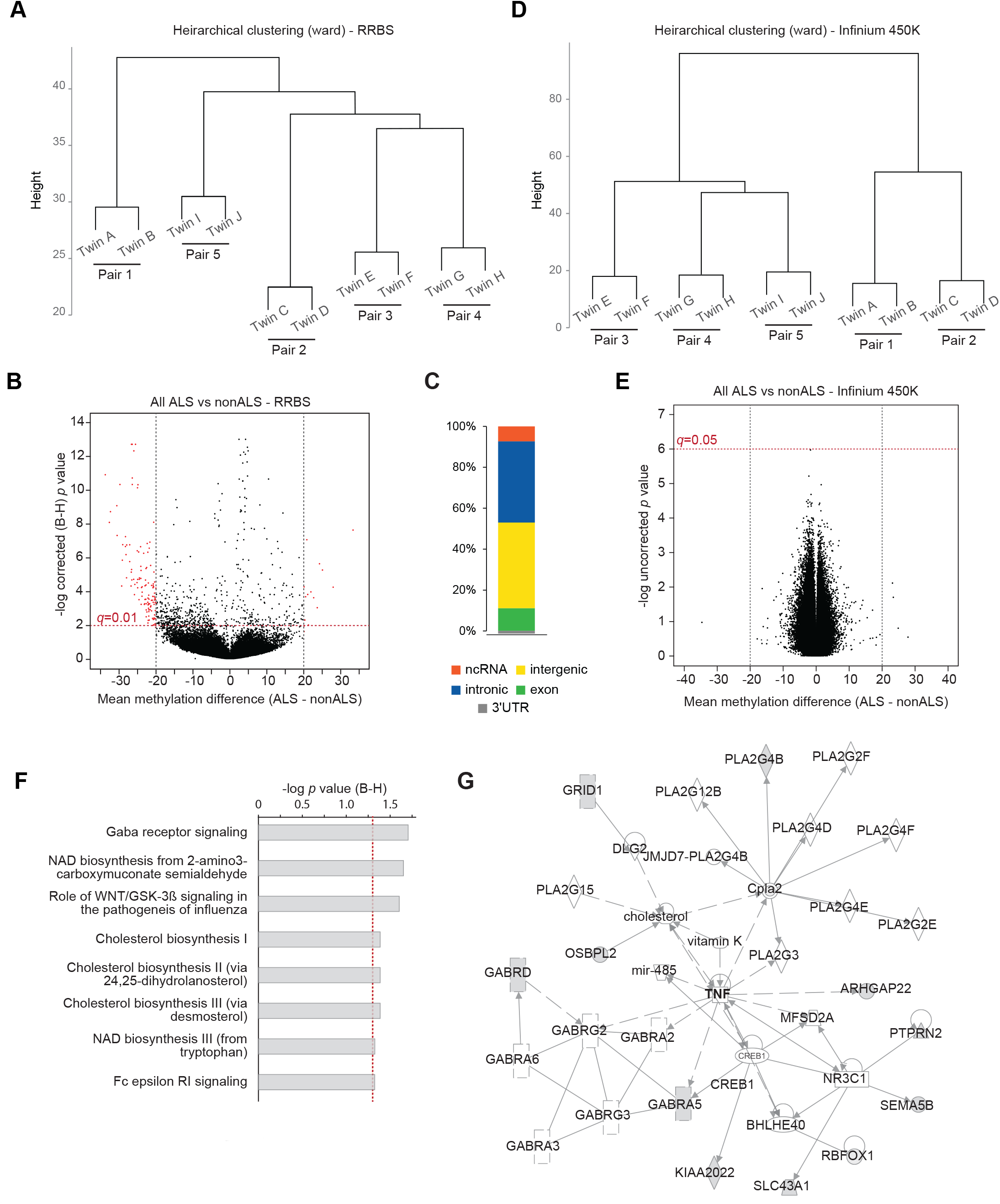
**RRBS reveals commonly differentially methylated CpGs in ALS cases that cluster in GABA receptor KIAA2022 SLC43A1 signaling. (A)** Dendrogram showing results of unsupervised hierarchical clustering of RRBS data at 10x coverage. **(B)** Volcano plot showing mean methylation difference between ALS cases and controls (x-axis) *vs*. -log corrected *p* values (y-axis) for CpG sites present in all RRBS libraries. Sites called as differentially methylated (at 20x coverage, 20% difference, *q*<0.01) are in red. **(C)** Genomic annotation of sites called as differentially methylated in (B). **(D)** Dendrogram showing results of unsupervised hierarchical clustering of Infinium 450K data. **(E)** Volcano plot showing mean methylation difference between ALS cases and controls (x-axis) *vs*. -log uncorrected *p* values (y-axis) for all CpG sites present on the 450K array. **(F)** Top canonical pathways represented by the genes harbouring differentially methylated cytosines between ALS cases and controls. **(G)** IPA network related to GABR signaling; genes harbouring differentially methylated cytosines are shaded grey.

None of the common DMCs identified by the RRBS case-control analysis exhibited changes consistent with a germline event (i.e. affecting most or all cells). On average the differences between cases and controls were ±25%, and while mosaicism for a germline change cannot be ruled out in this study of a single tissue, it is more likely that these modest changes indicate common somatic changes in ALS-affected individuals that are consequent to their disease. Ingenuity Pathway Analysis [29] (IPA) of the genes harbouring DMCs (*n*=74) revealed enrichment for several pathways, the most significantly enriched being ‘GABA receptor signalling’ (**Fig 2F**). IPA also identified four gene networks in which the affected genes function (**Fig S1**). The network containing genes involved in GABA signalling, shown in **Fig 2G**, centred around TNF. The other three pathways (two headed by cancer, and one by lipid metabolism) (**Fig S1**) have no obvious pathogenetic link to ALS, but since so little is known about the cause of ALS these networks warrant further investigation.

### Outlier analysis of RRBS data reveals characteristic epigenetic differences between ALS affected and unaffected twins

While the RRBS case-control analyses revealed interesting changes common to all twins, the necessary grouping of individuals for analysis means large changes of potential biological significance in only one or two ALS-affected individuals would be lost to statistical analysis. The ‘power of the twin’ would also be lost; this is particularly relevant in epigenetic studies, 155 where underlying DNA sequence can influence or even determine epigenetic state [30]. Given the clinical and genetic heterogeneity of ALS, the pathogenesis of motor neuron loss may be distinct in each affected twin. RRBS methylation patterns were therefore compared between each affected and unaffected individual in co-twin analyses.

We began by performing a Pearson's correlation of methylation levels between co-twins. Co-twin CpG methylation was highly correlated overall (r=0.978, range 0.972–0.982), and showed a generally bimodal distribution with most sites being either heavily methylated or largely unmethylated (**Fig 3A**). CpG sites present at >20x coverage in both twins within a pair were considered for further analysis. Those CpGs ≥5 residuals from the expected value from a linear model of all sites were called as methylation ‘outliers’ (**Fig 3B**). The minimum magnitude of difference in methylation at outliers between co-twins at this stringent cut-off was ~40%. Using this approach we identified more than 1,000 methylation outliers in each twin pair (**Fig 3C**; **Table S3**). Although there was a preponderance for methylation outliers to be hypomethylated in the ALS twins relative to the non-ALS twins, whole genome levels of 5-methylcytosine as measured by liquid chromatography-tandem mass spectrometry (LC-MS/MS) 170 did not differ between affected and unaffected individuals (**Fig 3D**), as has been previously suggested for ALS [31].

**Fig 3.**
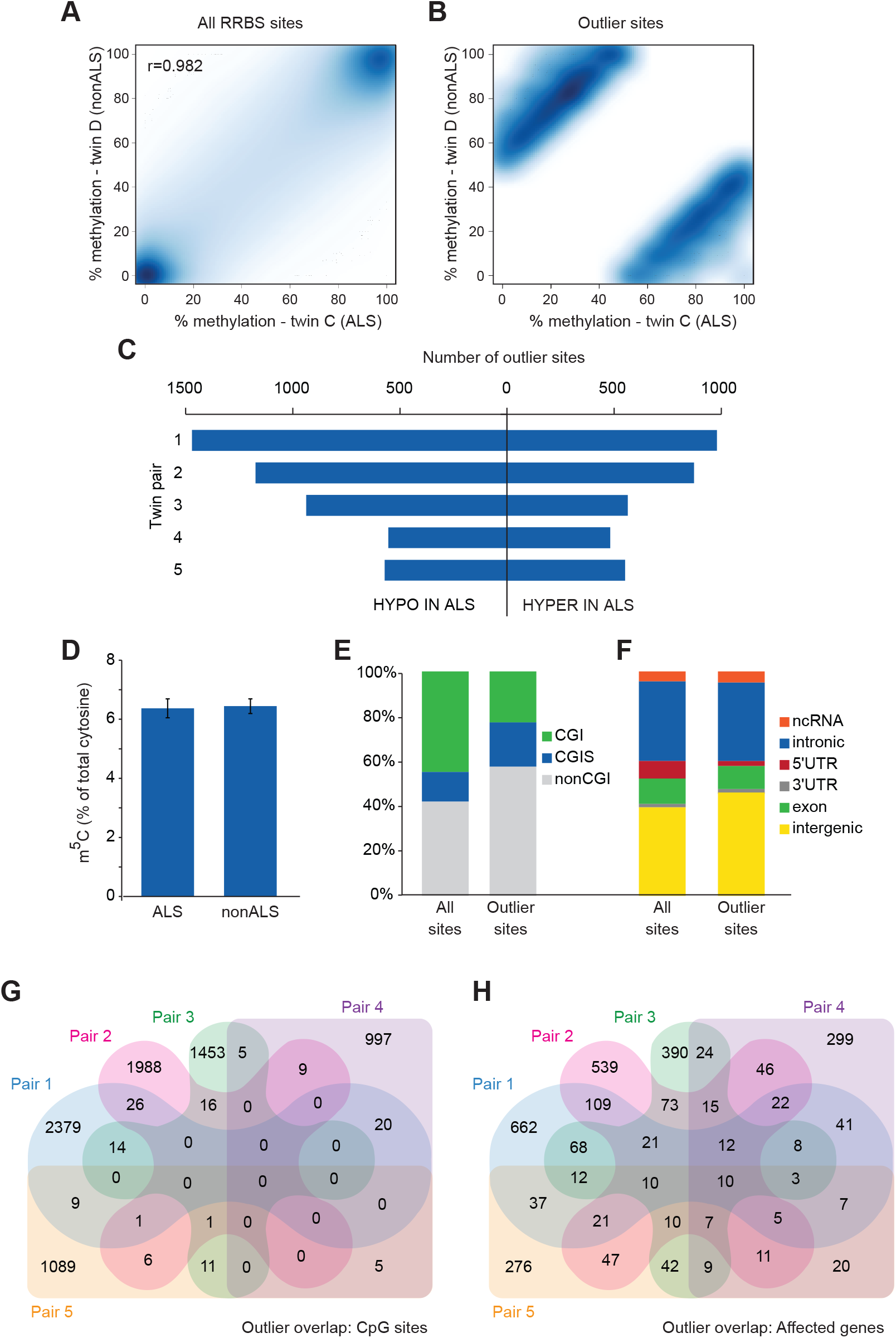
**Thousands of CpG sites show great discordance in methylation between ALS discordant co-twins. (A)** Smoothed correlation heatmap of all RRBS sites at 20x coverage in a representative twin pair (pair 2). **(B)** Smoothed correlation heatmap as in (A) showing only outlier sites ≥5 residuals from the linear model. **(C)** Bar graph showing the number of outliers defined by residuals in each twin pair. **(D)** Bar graph showing the total 5-methylcytosine content of peripheral blood DNA in ALS and nonALS individuals as measured by LC-MS/MS; error bars represent SEM. **(E,F)** Annotations for all RRBS sites and outlier sites for CpG islands **(E)** and genomic location **(F)**. **(G,H)** Venn diagrams showing overlaps among twin pairs for individual CpG outliers **(G)** and genes harbouring outliers **(H)**.

Genomic annotation of the outliers showed that, relative to all sites captured by RRBS, outlier sites were less likely to be in a CpG island (**Fig 3E**). Like the common DMCs identified by methylKit, outlier sites were predominantly in intronic and intergenic regions (**Fig 3F**). The majority of outlier CpGs were idiosyncratic to a twin pair, with little overlap among the twin pairs (**Fig 3G**). But when considering the genes harbouring the outlier CpG sites, the overlap among twins was greater, with ten genes (*ABR*, *NCOR2*, *SORCS2*, *HDAC4*, *SHANK2*, *RBFOX3*, *RXRA*, *MAD1L1*, *PTPRN2*, *GRIN1*) harbouring one or more methylation outliers in all five twin pairs (**Fig 3H**). Despite this overlap at the gene level, at least half of the affected genes were unique to a twin pair.

### ALS methylation outliers cluster in disease-relevant ontologies and pathways

We next took the genomic coordinates of the outlier CpGs and used the Genomic Regions Enrichment of Annotations Tool (GREAT) [32] to identify the ontologies of the sets of outliers for each twin pair. The molecular functions overrepresented by the outliers had one ontology in common across all twin pairs, ‘sequence specific DNA binding’ (**Table 2**). This is not disease-specific, but suggests that genes encoding transcription factors are susceptible to varying in epigenotype between identical genotypes. The significantly enriched biological functions revealed a large number of associated ontologies (**Table S4**), many of which may be relevant to disease. With the exception of twin pair 2, outliers of all twin pairs exhibited enriched biological functions that cluster in neurobiological pathways, including spinal cord and neuron development and differentiation (**Table 3**). Cellular compartment ontologies of the outliers were significantly enriched in three of the five twin pairs, all of which share a ‘Golgi lumen’ compartment enrichment (**Table 4)**. Golgi fragmentation is a well-recognised early event in multiple *in vitro* and animal models of ALS [33].

IPA analysis of the genes harbouring methylation outliers produced a set of top canonical pathways for each twin pair (**Table S5**). Cross-comparison of enriched pathways across all twin pairs revealed many significantly enriched pathways in common between two or more twin pairs (**Fig 4**). Most striking were the commonalities among neurobiological pathways, including pathways such as synaptic long-term potentiation. Taken together with the ontology analysis, this suggests that many methylation outliers represent an epigenetic signature of ALS in peripheral blood.

**Fig 4.**
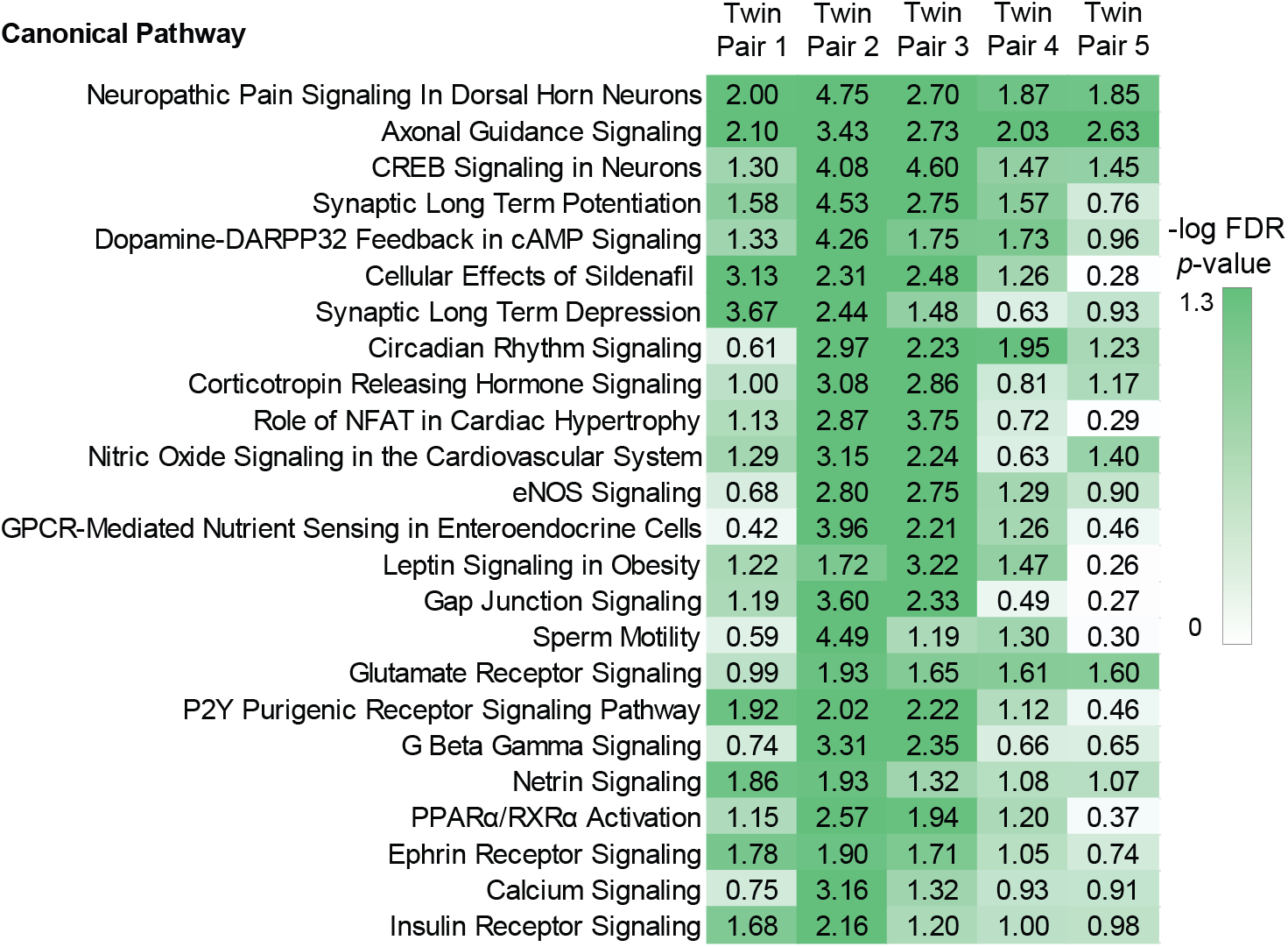
**Many canonical pathways associated with ALS methylation outliers are common among twin pairs.** Pseudoheatmap showing top canonical pathways common to more than one twin pair; -log *p* >1.3 signifies significant enrichment.

**Table 2.** Molecular functions associated with ALS methylation outliers

| **Molecular Function** | **Raw P-Value** | **FDR Q-Value** | **Fold Enrichment** |
| --- | --- | --- | --- |
| **Twin Pair 1** |  |  |  |
| sequence-specific DNA binding | 4.14E-19 | 9.55E-17 | 2.16 |
| **Twin Pair 2** |  |  |  |
| sequence-specific DNA binding transcription factor activity | 2.17E-23 | 6.66E-21 | 2.16 |
| nucleic acid binding transcription factor activity | 2.58E-23 | 7.32E-21 | 2.16 |
| sequence-specific DNA binding | 7.76E-22 | 2.05E-19 | 2.39 |
| regulatory region DNA binding | 1.51E-13 | 2.06E-11 | 2.45 |
| **Twin Pair 3** |  |  |  |
| sequence-specific DNA binding | 7.01E-14 | 1.08E-11 | 2.25 |
| tetrahydrobiopterin binding | 5.79E-07 | 3.45E-05 | 21.05 |
| **Twin Pair 4** |  |  |  |
| DNA binding | 3.09E-25 | 2.28E-22 | 2.18 |
| sequence-specific DNA binding | 6.68E-18 | 3.08E-15 | 2.81 |
| sequence-specific DNA binding transcription factor activity | 2.88E-15 | 1.18E-12 | 2.31 |
| nucleic acid binding transcription factor activity | 3.18E-15 | 1.17E-12 | 2.31 |
| regulatory region DNA binding | 5.65E-13 | 1.49E-10 | 3.10 |
| transcription regulatory region DNA binding | 2.65E-12 | 6.52E-10 | 3.05 |
| transcription regulatory region sequence-specific DNA binding | 7.76E-07 | 7.15E-05 | 3.09 |
| **Twin Pair 5** |  |  |  |
| transmembrane transporter activity | 6.56E-11 | 3.02E-08 | 2.11 |
| sequence-specific DNA binding | 7.12E-11 | 2.92E-08 | 2.25 |
| substrate-specific transmembrane transporter activity | 8.30E-11 | 3.06E-08 | 2.17 |
| ion transmembrane transporter activity | 1.18E-10 | 3.96E-08 | 2.20 |
| extracellular matrix structural constituent | 3.83E-08 | 6.42E-06 | 4.57 |
| cation channel activity | 6.66E-07 | 7.23E-05 | 2.54 |

**Table 3.** Neurobiological^a^ processes associated 204 with ALS methylation outliers

| Biological process | Raw P-Value | FDR Q-Val | Fold Enrichment |
| --- | --- | --- | --- |
| **Twin Pair 1** |  |  |  |
| cell differentiation in spinal cord | 4.46E-10 | 1.59E-08 | 4.00 |
| spinal cord development | 2.22E-09 | 7.30E-08 | 3.18 |
| dorsal spinal cord development | 5.14E-09 | 1.61E-07 | 6.25 |
| spinal cord association neuron differentiation | 5.10E-06 | 8.95E-05 | 5.79 |
| **Twin Pair 3** |  |  |  |
| generation of neurons | 1.21E-23 | 2.21E-21 | 2.21 |
| neurogenesis | 3.95E-23 | 6.65E-21 | 2.16 |
| neuron differentiation | 2.33E-20 | 3.29E-18 | 2.28 |
| neuron development | 1.08E-14 | 9.83E-13 | 2.18 |
| neuron projection morphogenesis | 5.78E-12 | 3.55E-10 | 2.25 |
| central nervous system development | 9.64E-12 | 5.78E-10 | 2.02 |
| **Twin Pair 4** |  |  |  |
| central nervous system development | 8.78E-14 | 1.01E-11 | 2.39 |
| brain development | 6.27E-12 | 5.85E-10 | 2.49 |
| **Twin Pair 5** |  |  |  |
| nervous system development | 9.48E-20 | 3.96E-17 | 2.00 |
| generation of neurons | 1.63E-13 | 3.09E-11 | 2.01 |
| regulation of nervous system development | 3.74E-08 | 3.15E-06 | 2.19 |
| regulation of neurogenesis | 1.61E-07 | 1.19E-05 | 2.20 |
| negative regulation of neuron differentiation | 5.73E-06 | 2.74E-04 | 4.43 |

<sup>a</sup> A full list of all enriched biological process can be found in Table S4

**Table 4.** Cellular components associated with 207 ALS methylation outliers

| Cellular Component | Raw P-Value | FDR Q-Val | Fold Enrichment |
| --- | --- | --- | --- |
| **Twin Pair 1** |  |  |  |
| Golgi lumen | 6.55E-16 | 2.37E-14 | 4.85 |
| **Twin Pair 4** |  |  |  |
| extracellular matrix | 2.18E-12 | 7.06E-11 | 2.24 |
| proteinaceous extracellular matrix | 4.28E-11 | 1.32E-09 | 2.27 |
| anchoring junction | 3.77E-10 | 1.04E-08 | 2.62 |
| adherens junction | 1.83E-09 | 4.82E-08 | 2.61 |
| Golgi lumen | 8.03E-09 | 1.92E-07 | 3.82 |
| cell-cell adherens junction | 4.23E-05 | 0.000547 | 3.33 |
| **Twin Pair 5** |  |  |  |
| extracellular matrix part | 1.74E-05 | 4.78E-04 | 2.53 |
| Golgi lumen | 5.22E-05 | 1.16E-03 | 3.62 |
| voltage-gated potassium channel complex | 7.20E-05 | 1.49E-03 | 3.51 |
| cation channel complex | 1.68E-04 | 2.95E-03 | 2.57 |
| ion channel complex | 1.78E-04 | 3.04E-03 | 2.16 |

## Discussion

We have taken advantage of the genetic and early environmental similarity of identical twins discordant for ALS to gain insight into the nature and extent of epigenetic changes in this disease. Together our findings demonstrate that ALS has epigenetic signatures in peripheral blood DNA that could potentially be exploited as biomarkers of disease. Our findings are consistent with widespread disruptions to epigenetic patterns in ALS that either underlie disease etiology, or represent changes consequent to pathology.

Familial ALS is genetically heterogeneous, but clinically very similar to SALS, which prompted us to use our data to first examine methylation at genes known to be mutated in familial ALS. Germline epimutation, characterised by soma-wide aberrant silencing of a gene, can phenocopy a genetic mutation [34], and is usually associated with dense hypermethylation at the promoter of the affected gene. However none of the individuals exhibited any aberrant methylation at known ALS gene promoters in their peripheral blood. This finding does not necessarily preclude an inborn epigenetic defect as the basis for an affected twin's predisposition to ALS, but it excludes this possibility at known ALS genes.

Unbiased case-control analyses are designed to detect commonalities between groups. It is of particular interest that our RRBS analyses revealed affected twin-concordant methylation changes at genes that cluster in GABA receptor signalling. Cortical hyperexcitability is one of the earliest identifiable changes in patients with ALS, caused at least in part by degeneration of inhibitory cortical circuits and reduced cortical GABA levels [35, 36]. Given that ALS is such a heterogeneous disease [37], these epigenetic changes common to all our ALS affected twins could be to secondary to the many pathogenetic pathways found in ALS, rather than being causally related to the disease. If so, these changes hold the potential to be exploited as blood-based biomarkers for an early diagnosis of ALS.

When considering methylation differences between twins we found a considerable number of differences of large magnitude and defined these as ‘methylation outliers’. Based on the magnitude of difference in methylation between co-twins at outliers and the stringent parameters we used to identify them, it is unlikely that these outliers merely reflect experimental noise. We do not expect, however, that all methylation outliers between co-twins will be representative of ALS discordance, since many differences may reflect or underlie other phenotypic discordances, 239 or individual exposure to environmental factors [12]. For example, one of our individuals was a smoker at the time of sample collection and her co-twin was not; in this pair we were able to identify the expected difference in methylation levels at an intronic CpG in the *AHRR* gene, 242 known to robustly associated with active smoking [38] (Fig S2). This particular difference fell just under our outlier threshold of ≥5 residuals, but given that twin pairs carry thousands of outlier sites of greater magnitude than this, at least some of them will be expected to reflect the discordance for ALS, a supposition supported by the gene ontology and pathway analyses of outliers. Genome-wide analyses of outliers identified in healthy twins (performed in a similar manner [12]) revealed between-twin differences that cluster largely in ontologies related to the tissue being examined; between-twin DMCs in adipose tissue clustered in functions related to lipid metabolism while peripheral blood DMCs clustered in haematological functions [12].

The thousands of outlier sites we identified in each twin pair showed only a modest overlap in genes affected, but all five twin pairs harboured outliers in ten common genes. Three of these genes have previously been implicated in ALS: *SORCS2*, *RXRA*, and *HDAC4*, which have prominent roles in inflammation and epigenetic regulation [39–41]. *GRIN1*, another of the ten common genes, encodes a subunit of the glutamate NMDA receptor, the major mediator of excitotoxicity; splicing of *GRIN1* requires the RNA binding protein TAF15, another molecule implicated in ALS [42]. The remaining genes, including *ABR*, *SHANK2*, *RBFOX3* and *PTPRN2* have no obvious link to ALS, but are notable for being highly expressed in the central nervous system. The genes which are affected in all our cases could be considered candidates in follow-up studies of larger SALS cohorts.

The overlap in functional pathways and networks associated with ALS methylation outliers was the most striking finding of this study. Neurobiological functions or pathways relevant to ALS were overrepresented in every twin pair, even with the modest lack of gene overlap, and more importantly, with the tissue that was examined (white blood cells, not CNS). We were not able to adjust for blood cell composition but such differences, if present, would not be expected to result in enrichment for neurobiological-related ontologies. Perturbed neuro-related pathways in non-affected tissue might reflect different routes to the common endpoint of ALS in each affected twin; these could potentially be germline epigenetic changes that have predisposed to ALS, but we are unable to establish this as other normal tissues were not available for analysis. On the other hand, it is equally plausible (if not more likely) that the idiosyncratic CpG outliers in affected twins are representative of different environmental exposures, some of which have contributed to ALS susceptibility. Assessing larger cohorts of sporadic ALS for the presence of the outliers identified in this study may yield greater insight into their role in ALS.

A noteworthy finding of this study is that the differences we identified with RRBS could not be detected with the 450K array, because the majority of ALS methylation outliers we found are not represented on the array. While the 450K array has been a popular method for epigenetic epidemiology due to its low cost and ease of analysis, our results show that the representative set of CpGs on the array are less than optimal in capturing the extent of epigenetic variation in ALS. RRBS captures only around 1% of the genome (although enriched for CpGs), but with the increasing affordability of high-throughput sequencing, whole genome bisulfite sequencing (WGBS) of large cohorts will soon be become feasible. Our results suggest that future WGBS studies will be required to capture the full extent of epigenetic discordance among identical twins with discordant disease phenotypes.

## Materials and Methods

### Ethics statement

Informed written consent was obtained from each individual for their DNA to be used in the study protocol ‘Looking for the Causes of MND’, approved by the Sydney South West Area Health Service Human Research Ethics Committee (no. X11-0383 & HREC/11/RPAH/601).

### Participants

Five individuals with a diagnosis of SALS and their unaffected monozygotic twin siblings were involved in this study. The diagnosis of SALS was made by a neurologist, with four having classical ALS (with upper and motor neuron signs) and one with the progressive muscular atrophy (PMA) variant (with lower motor neurons signs only). Autopsy neuropathological confirmation of the diagnosis was available for one patient with classical ALS and one with the PMA variant. No twin had a family history of ALS. All affected and unaffected co-twins donated blood samples to the Australian Motor Neuron Disease DNA Bank and completed a detailed demographic and environmental exposure questionnaire. Epidemiological and clinical differences between the co-twins are shown in Table 1. Venous blood samples were taken from an antecubital vein at the same time in each twin pair. DNA was extracted from white blood cells using the QIAmp blood kit (Qiagen) and stored at -20°C until used.

### Total 5-methylcytosine (5mc) content

Total 5mc content of each DNA sample was analysed by liquid chromatography-mass spectrometry (LC-MS/MS). Approximately 1 μg of genomic DNA was used in hydrolysis using DNA Degradase Plus (Zymo). The reaction mixture was incubated at 37°C for two hours to ensure complete digestion prior to LC-MS/MS, as described previously [43].

### Reduced representation bisulfite sequencing (RRBS)

Indexed RRBS libraries were prepared from 1μg of *Msp*I-digested genomic DNA essentially as described [24], and sequenced in multiplex on the Illumina HiSeq 2000. Resulting fastq files were trimmed with cutadapt v1.3. Trimmed reads were aligned to the human reference genome (hg19) using Bismark v0.10.0 [44] paired with Bowtie v1 [45] with default parameters with methylation calling by Bismark-methylation-extractor. Output files were reformatted for direct input into methylKit using a custom script.

### RRBS case-control analysis

Differentially methylated CpG sites between all cases and controls were identified using the 315 Bioconductor R package methylKit [27] with filter settings of ≥20X coverage ≥20% methylation difference, and *q* value of 0.01.

### Outlier analysis

Linear models were established using R for each twin pair using methylation calls for CpG sites in common to co-twins with ≥20x coverage. Outlier CpG sites were defined as those ≥5 residuals from the predicted value from the linear model. Genomic coordinates for outlier sites for each twin pair were analysed with the gene ontology software GREAT [32]. Genes harbouring outliers were analysed further by Ingenuity Pathway Analysis (http://www.ingenuity.com/).

### Illumina Infinium 450K arrays

Infinium 450K arrays were performed on each sample by the Australian Genome Research Facility (http://www.agrf.org.au/). Resultant data were analysed using the Bioconductor package minfi[28] using SWAN normalisation. Only probes with a detection value of *p* value <0.01 were included in analysis.

### Data availability

All raw data generated by this study (RRBS and 450K array) have been deposited at the NCBI Gene Expression Omnibus under Accession Number GSE89474.

## Acknowledgements

We thank ALS patients and their twin siblings for donating DNA samples, treating neurologists for supplying clinical information, MND Associations in all Australian states for assisting with sample collections, and Roland Stocker, Suzy Hur, and Ghassan Maghzal for performing the LC-MS/MS analysis.

## References

1. Kiernan MC, Vucic S, Cheah BC, Turner MR, Eisen A, Hardiman O, et al. Amyotrophic lateral sclerosis. Lancet. 2011;377(9769):942–55. Epub 2011/02/08. doi:S0140-6736(10)61156-7 [pii]10.1016/S0140-6736(10)61156-7. PubMed PMID: 21296405; PubMed Central PMCID: PMCALS.

2. Al-Chalabi A, Calvo A, Chio A, Colville S, Ellis CM, Hardiman O, et al. Analysis of amyotrophic lateral sclerosis as a multistep process: a population-based modelling study. Lancet Neurol. 2014;13(11):1108–13. doi:Doi 10.1016/S1474-4422(14)70219-4. PubMed PMID: WOS:000343783900018.

3. Steinberg KM, Yu B, Koboldt DC, Mardis ER, Pamphlett R. Exome sequencing of case-unaffected-parents trios reveals recessive and de novo genetic variants in sporadic ALS. Scientific reports. 2015;5:9124. doi:10.1038/srep09124. PubMed PMID: 25773295.

4. Couthouis J, Raphael AR, Daneshjou R, Gitler AD. Targeted exon capture and sequencing in sporadic amyotrophic lateral sclerosis. PLoS genetics. 2014;10(10):e1004704. doi:10.1371/journal.pgen.1004704. PubMed PMID: 25299611; PubMed Central PMCID: PMCPMC4191946.

5. Keller MF, Ferrucci L, Singleton AB, Tienari PJ, Laaksovirta H, Restagno G, et al. Genome-Wide Analysis of the Heritability of Amyotrophic Lateral Sclerosis. JAMA neurology. 2014. Epub 2014/07/16. doi:10.1001/jamaneurol.2014.1184. PubMed PMID: 25023141.

6. Belzil VV, Katzman RB, Petrucelli L. ALS and FTD: an epigenetic perspective. Acta Neuropathol. 2016. doi:10.1007/s00401-016-1587-4. PubMed PMID: 27282474.

7. Paez-Colasante X, Figueroa-Romero C, Sakowski SA, Goutman SA, Feldman EL. Amyotrophic lateral sclerosis: mechanisms and therapeutics in the epigenomic era. Nat Rev Neurol. 2015;11(5):266–79. doi:10.1038/nrneurol.2015.57. PubMed PMID: 25896087.

8. Oates N, Pamphlett R. An epigenetic analysis of SOD1 and VEGF in ALS. Amyotroph Lateral Scler. 2007;8(2):83–6. doi:10.1080/17482960601149160. PubMed PMID: 17453634.

9. Morahan JM, Yu B, Trent RJ, Pamphlett R. Are metallothionein genes silenced in ALS? Toxicology letters. 2007;168(1):83–7. PubMed PMID: 17156946.

10. Morahan JM, Yu B, Trent RJ, Pamphlett R. A genome-wide analysis of brain DNA methylation identifies new candidate genes for sporadic amyotrophic lateral sclerosis. Amyotroph Lateral Scler. 2009;10(5-6):418–29. doi:10.3109/17482960802635397. PubMed PMID: 19922134.

11. Al-Chalabi A, Kwak S, Mehler M, Rouleau G, Siddique T, Strong M, et al. Genetic and epigenetic studies of amyotrophic lateral sclerosis. Amyotroph Lateral Scler Frontotemporal Degener. 2013;14 Suppl 1:44–52. Epub 2013/05/25. doi:10.3109/21678421.2013.778571. PubMed PMID: 23678879.

12. Busche S, Shao X, Caron M, Kwan T, Allum F, Cheung WA, et al. Population whole-genome bisulfite sequencing across two tissues highlights the environment as the principal source of human methylome variation. Genome Biol. 2015;16:290. doi:10.1186/s13059-015-0856-1. PubMed PMID: 26699896; PubMed Central PMCID: PMCPMC4699357.

13. Ladd-Acosta C, Fallin MD. The role of epigenetics in genetic and environmental epidemiology. Epigenomics. 2016;8(2):271–83. doi:10.2217/epi.15.102. PubMed PMID: 26505319.

14. Graham AJ, Macdonald AM, Hawkes CH. British motor neuron disease twin study. Journal of neurology, neurosurgery, and psychiatry. 1997;62(6):562–9. Epub 1997/06/01. PubMed PMID: 9219739; PubMed Central PMCID: PMC1074137.

15. Dellefave L, Bangash MA, Siddique T. Pairwise concordance rates are similar in monozygotic and dizygotic twins for amyotrophic lateral sclerosis. Amyotroph Lateral Scler Other Motor Neuron Disord. 2003;4 Suppl 1:47. PubMed Central PMCID: PMCALS.

16. Al-Chalabi A, Fang F, Hanby MF, Leigh PN, Shaw CE, Ye W, et al. An estimate of amyotrophic lateral sclerosis heritability using twin data. Journal of neurology, neurosurgery, and psychiatry. 2010;81(12):1324–6. Epub 2010/09/24. doi:10.1136/jnnp.2010.207464. PubMed PMID: 20861059; PubMed Central PMCID: PMC2988617.

17. Kaminsky ZA, Tang T, Wang SC, Ptak C, Oh GH, Wong AH, et al. DNA methylation profiles in monozygotic and dizygotic twins. Nat Genet. 2009;41(2):240–5. doi:10.1038/ng.286. PubMed PMID: 19151718.

18. Gervin K, Vigeland MD, Mattingsdal M, Hammero M, Nygard H, Olsen AO, et al. DNA methylation and gene expression changes in monozygotic twins discordant for psoriasis: identification of epigenetically dysregulated genes. PLoS genetics. 2012;8(1):e1002454. doi:10.1371/journal.pgen.1002454. PubMed PMID: 22291603; PubMed Central PMCID: PMC3262011.

19. Vogt J, Kohlhase J, Morlot S, Kluwe L, Mautner VF, Cooper DN, et al. Monozygotic twins discordant for neurofibromatosis type 1 due to a postzygotic NF1 gene mutation. Human mutation. 2011;32(6):E2134–47. Epub 2011/05/28. doi:10.1002/humu.21476. PubMed PMID: 21618341.

20. Robertson SP, Jenkins ZA, Morgan T, Ades L, Aftimos S, Boute O, et al. Frontometaphyseal dysplasia: mutations in FLNA and phenotypic diversity. American journal of medical genetics Part A. 2006;140(16):1726–36. doi:10.1002/ajmg.a.31322. PubMed PMID: 16835913.

21. Pamphlett R, Cheong PL, Trent RJ, Yu B. Can ALS-associated C9orf72 repeat expansions be diagnosed on a blood DNA test alone? PLoS One. 2013;8(7):e70007. doi:10.1371/journal.pone.0070007. PubMed PMID: 23894576; PubMed Central PMCID: PMCPMC3716700.

22. Meltz Steinberg K, Nicholas TJ, Koboldt DC, Yu B, Mardis E, Pamphlett R. Whole genome analyses reveal no pathogenetic single nucleotide or structural differences between monozygotic twins discordant for amyotrophic lateral sclerosis. Amyotroph Lateral Scler Frontotemporal Degener. 2015;16(5-6):385–92. doi:10.3109/21678421.2015.1040029. PubMed PMID: 25960086.

23. Sandoval J, Heyn H, Moran S, Serra-Musach J, Pujana MA, Bibikova M, et al. Validation of a DNA methylation microarray for 450,000 CpG sites in the human genome. Epigenetics. 2011;6(6):692–702. PubMed PMID: 21593595.

24. Gu H, Smith ZD, Bock C, Boyle P, Gnirke A, Meissner A. Preparation of reduced representation bisulfite sequencing libraries for genome-scale DNA methylation profiling. Nat Protoc. 2011;6(4):468–81. doi:10.1038/nprot.2010.190. PubMed PMID: 21412275.

25. Renton AE, Chio A, Traynor BJ. State of play in amyotrophic lateral sclerosis genetics. Nat Neurosci. 2014;17(1):17–23. doi:10.1038/nn.3584. PubMed PMID: 24369373; PubMed Central PMCID: PMCPMC4544832.

26. McRae AF, Powell JE, Henders AK, Bowdler L, Hemani G, Shah S, et al. Contribution of genetic variation to transgenerational inheritance of DNA methylation. Genome Biol. 2014;15(5):R73. doi:10.1186/gb-2014-15-5-r73. PubMed PMID: 24887635; PubMed Central PMCID: PMCPMC4072933.

27. Akalin A, Kormaksson M, Li S, Garrett-Bakelman FE, Figueroa ME, Melnick A, et al. methylKit: a comprehensive R package for the analysis of genome-wide DNA methylation profiles. Genome Biol. 2012;13(10):R87. doi:10.1186/gb-2012-13-10-r87. PubMed PMID: 23034086; PubMed Central PMCID: PMCPMC3491415.

28. Aryee MJ, Jaffe AE, Corrada-Bravo H, Ladd-Acosta C, Feinberg AP, Hansen KD, et al. Minfi: a flexible and comprehensive Bioconductor package for the analysis of Infinium DNA methylation microarrays. Bioinformatics. 2014;30(10):1363–9. doi:10.1093/bioinformatics/btu049. PubMed PMID: 24478339; PubMed Central PMCID: PMCPMC4016708.

29. www.qiagen.com/ingenuity.

30. Richards EJ., Inherited epigenetic variation–revisiting soft inheritance. Nat Rev Genet. 2006;7(5):395–401. doi:10.1038/nrg1834. PubMed PMID: 16534512.

31. Tremolizzo L, Messina P, Conti E, Sala G, Cecchi M, Airoldi L, et al. Whole-blood global DNA methylation is increased in amyotrophic lateral sclerosis independently of age of onset. Amyotroph Lateral Scler Frontotemporal Degener. 2014;15(1-2):98–105. doi:10.3109/21678421.2013.851247. PubMed PMID: 24224837.

32. McLean CY, Bristor D, Hiller M, Clarke SL, Schaar BT, Lowe CB, et al. GREAT improves functional interpretation of cis-regulatory regions. Nat Biotechnol. 2010;28(5):495–501. doi:10.1038/nbt.1630. PubMed PMID: 20436461; PubMed Central PMCID: PMCPMC4840234.

33. Haase G, Rabouille C. Golgi Fragmentation in ALS Motor Neurons. New Mechanisms Targeting Microtubules, Tethers, and Transport Vesicles. Front Neurosci. 2015;9:448. doi:10.3389/fnins.2015.00448. PubMed PMID: 26696811; PubMed Central PMCID: PMCPMC4672084.

34. Martin DI, Ward R, Suter CM. Germline epimutation: A basis for epigenetic disease in humans. Ann N Y Acad Sci. 2005;1054:68–77. doi:10.1196/annals.1345.009. PubMed PMID: 16339653.

35. Vucic S, Cheah BC, Kiernan MC. Defining the mechanisms that underlie cortical hyperexcitability in amyotrophic lateral sclerosis. Exp Neurol. 2009;220(1):177–82. doi:10.1016/j.expneurol.2009.08.017. PubMed PMID: 19716820.

36. Foerster BR, Callaghan BC, Petrou M, Edden RA, Chenevert TL, Feldman EL. Decreased motor cortex gamma-aminobutyric acid in amyotrophic lateral sclerosis. Neurology. 2012;78(20):1596–600. doi:10.1212/WNL.0b013e3182563b57. PubMed PMID: 22517106; PubMed Central PMCID: PMCPMC3348851.

37. Robberecht W, Philips T. The changing scene of amyotrophic lateral sclerosis. Nat Rev Neurosci. 2013;14(4):248–64. doi:10.1038/nrn3430. PubMed PMID: 23463272.

38. Gao X, Jia M, Zhang Y, Breitling LP, Brenner H. DNA methylation changes of whole blood cells in response to active smoking exposure in adults: a systematic review of DNA methylation studies. Clin Epigenetics. 2015;7:113. doi:10.1186/s13148-015-0148-3. PubMed PMID: 26478754; PubMed Central PMCID: PMCPMC4609112.

39. Mori F, Miki Y, Tanji K, Kakita A, Takahashi H, Utsumi J, et al. Sortilin-related receptor CNS expressed 2 (SorCS2) is localized to Bunina bodies in amyotrophic lateral sclerosis. Neurosci Lett. 2015;608:6–11. doi:10.1016/j.neulet.2015.09.030. PubMed PMID: 26420026.

40. Brohawn DG, O'Brien LC, Bennett JP Jr. RNAseq Analyses Identify Tumor Necrosis Factor-Mediated Inflammation as a Major Abnormality in ALS Spinal Cord. PLoS One. 2016;11(8):e0160520. doi:10.1371/journal.pone.0160520. PubMed PMID: 27487029; PubMed Central PMCID: PMCPMC4972368.

41. Bruneteau G, Simonet T, Bauche S, Mandjee N, Malfatti E, Girard E, et al. Muscle histone deacetylase 4 upregulation in amyotrophic lateral sclerosis: potential role in reinnervation ability and disease progression. Brain. 2013;136(Pt 8):2359–68. doi:10.1093/brain/awt164. PubMed PMID: 23824486.

42. Ibrahim F, Maragkakis M, Alexiou P, Maronski MA, Dichter MA, Mourelatos Z. Identification of in vivo, conserved, TAF15 RNA binding sites reveals the impact of TAF15 on the neuronal transcriptome. Cell Rep. 2013;3(2):301–8. doi:10.1016/j.celrep.2013.01.021. PubMed PMID: 23416048; PubMed Central PMCID: PMCPMC3594071.

43. Le T, Kim KP, Fan G, Faull KF. A sensitive mass spectrometry method for simultaneous quantification of DNA methylation and hydroxymethylation levels in biological samples. Anal Biochem. 2011;412(2):203–9. doi:10.1016/j.ab.2011.01.026. PubMed PMID: 21272560; PubMed Central PMCID: PMCPMC3070205.

44. Krueger F, Andrews SR. Bismark: a flexible aligner and methylation caller for Bisulfite-Seq applications. Bioinformatics. 2011;27(11):1571–2. doi:10.1093/bioinformatics/btr167. PubMed PMID: 21493656; PubMed Central PMCID: PMC3102221.

45. Langmead B, Trapnell C, Pop M, Salzberg SL. Ultrafast and memory-efficient alignment of short DNA sequences to the human genome. Genome Biol. 2009;10(3):R25. doi:10.1186/gb-2009-10-3-r25. PubMed PMID: 19261174; PubMed Central PMCID: PMC2690996.

